# Evaluating the efficacy of multiple myeloma cell lines as models for patient tumors via transcriptomic correlation analysis

**DOI:** 10.1101/847368

**Authors:** Vishesh Sarin, Katharine Yu, Ian D. Ferguson, Olivia Gugliemini, Matthew A. Nix, Byron C. Hann, Marina Sirota, Arun P. Wiita

## Abstract

Multiple myeloma (MM) cell lines are routinely used to model the disease. However, a long-standing question is how well these cell lines truly represent tumor cells in patients. Here, we employ a recently-described method of transcriptional correlation profiling to compare similarity of 66 MM cell lines to 779 newly-diagnosed MM patient tumors. We found that individual MM lines differ significantly with respect to patient tumor representation, with median *R* ranging from 0.35-0.54. ANBL-6 was the “best” line, markedly exceeding all others (*p* < 2.2e-16). Notably, some widely-used cell lines (RPMI-8226, U-266) scored poorly in our patient similarity ranking (48 and 52 of 66, respectively). Lines cultured with interleukin-6 showed significantly improved correlations with patient tumor (*p* = 9.5e-4). When common MM genomic features were matched between cell lines and patients, only t(4;14) and t(14;16) led to increased transcriptional correlation. To demonstrate utility of our top-ranked line for preclinical studies, we showed that intravenously-implanted ANBL-6 proliferates in hematopoietic organs in immunocompromised mice. Overall, our large-scale quantitative correlation analysis, utilizing emerging datasets, provides a resource informing the MM community of cell lines that may be most reliable for modeling patient disease while also elucidating biological differences between cell lines and tumors.

## Introduction

The past 20 years have seen remarkable advances in multiple myeloma (MM) biology and therapy. Many of these discoveries originated with studies performed in MM cell lines. However, there are long-standing questions about the reliability of MM cell lines as models for true disease in patients. One major issue is that while the large majority of MM tumor cells reside within the bone marrow niche, essentially all MM cell lines have been derived from disease growing outside the bone marrow^1^. These patient cells of origin were either circulating in the bloodstream in the form of plasma cell leukemia or in effusions at other sites^2^. In both cases, these cells are expected to have lost reliance on the bone marrow microenvironment for proliferation. Intimate dependence on the marrow niche is a well-known hallmark of typical MM biology^3–5^. However, attempts to establish long-term culture of MM plasma cells isolated from purely marrow-localized disease have been largely unsuccessful^2^. Therefore, there is ample reason to presume that MM cell lines will carry different phenotypes from patient disease *in vivo*.

Despite these caveats, cell lines remain the workhorse of MM research. To mitigate these limitations, several groups have developed cell lines that remain dependent on interleukin-6 (IL-6) in culture media^1, 2, 6^. IL-6 is recognized as the most critical bone marrow microenvironment factor supporting MM tumor growth^7–9^. Therefore, these lines may recapitulate additional *in vivo* phenotypes lost in IL-6 independent lines. In parallel, many attempts have been made to match detected genomic alterations in cell lines, such as translocations or mutations, to specific experimental phenotypes, and then extrapolate these findings to patients with the same genomic lesions (for example, refs.^10–12^). However, it is unclear how generally the genotype-associated observations in cell lines truly relate to effects of those genotypes in patient tumor.

Taken together, significant questions remain about both the qualitative and quantitative differences between MM cell lines and patient tumors. Furthermore, it remains unclear whether specific cell lines are more representative of patient tumors than others. Here, we aim to address these questions. Our work extends from our recent study^13^, where we correlated RNA-seq data from 666 cell lines in the Cancer Cell Line Encyclopedia (CCLE) to each patient’s RNA-seq data in The Cancer Genome Atlas (TCGA), derived from 8,282 tumors, across 22 matching tumor types. The central hypothesis of this approach is that global gene expression patterns provide the most robust phenotypic representation of cellular biology. We specifically identified cell lines that showed greatly increased and decreased global transcriptomic correlations versus primary patient samples. Based on these results, we proposed that the cell lines used in the standard “NCI-60” preclinical panel should be replaced by a “TCGA-110-CL”, employing a cohort of lines with the most similarity to patient tumors. In parallel, others have also used transcriptional correlation profiling to suggest the best cell line models of metastatic breast cancer^14^ and hepatocellular carcinoma^15^, for example, demonstrating the widespread utility of this approach.

As the TCGA primarily includes data on solid tumors, our prior publication did not include MM. Fortunately, the Multiple Myeloma Research Foundation (MMRF) has addressed this gap in knowledge. The MMRF has sponsored a comprehensive transcriptomic resource of MM cell lines (www.keatslab.org) and MM patient tumors within the MMRF CoMMpass study (research.themmrf.org). Here, we employed our transcriptional correlation profiling approach to perform 51,414 individual correlations of cell lines vs. patient tumor. We confirmed that MM cell lines and patient tumors display broad transcriptomic differences. However, we did identify cell lines, in particular ANBL-6, that appear to be more representative of patient disease than others. In contrast, some widely used lines scored relatively poorly in our ranking of similarity to patients. We further characterized additional features to aid in increasing similarity of cell lines to patient tumor. Here, we provide a resource for cell line selection in MM research while also elucidating underlying biological signatures distinguishing cell lines and patient tumors.

## Materials and Methods

### Transcriptome and Mutational Analysis

See details of analysis in Supplementary Methods. Briefly, annotated read count data was obtained from Keats lab cell line (www.keatslab.org) and CoMMpass IA13 patient datasets (research.themmrf.org) and normalized via variance stabilizing transformation^16^. The top 5000 most variable genes were used for Spearman correlation analyses. Exome sequencing-based mutation data was similarly obtained from annotated datasets from these resources. Clinical subset analysis was performed as annotated for patients in CoMMpass. CoMMpass patient translocations were annotated as in ref.^17^

### ANBL-6 experiments

See details of analysis in Supplementary Methods. Briefly, ANBL-6 cell lines were stably transduced with a lentiviral construct stably expressed enhanced firefly luciferase and implanted into female 6-8 week old NOD *scid* gamma (NSG) mice. Tumor burden was monitored by bioluminescent imaging.

## Results

### Global correlation analysis reveals MM cell lines are not equal representations of patient tumors

We began by obtaining RNA-seq read count data for 66 MM cell lines (Keats lab resource) and CD138+ enriched tumor cells from 779 newly-diagnosed MM patients (CoMMpass release IA13). As in our prior study^13^, we normalized all reads using the upper-quartile method via Variance Stabilizing Transformation^16^ (see Methods). We note that all cell line and patient RNA sequencing libraries were prepared and analyzed in the same laboratory (Jonathan Keats lab at TGEN), reducing potential for artifacts when comparing samples generated from different groups.

As in our prior study, we focused our analysis on the top 5,000 most variable genes across samples expressed consistently at >1 counts per million, with the reasoning that these genes are most likely to be biologically informative for similarity assessment (see Supplementary Methods). A workflow for our analysis is shown in Fig. 1. Our primary analysis is performing a Spearman correlation across these 5,000 genes for each cell line versus each patient tumor sample, with the hypothesis that a perfect correlation (*R* = 1) means that a cell line is an exact representation of the patient tumor. We show individual correlation plots in Fig. 2 to provide examples of the 51,414 total correlations performed to generate our overall rankings in Fig. 3A.

**Figure 1.**
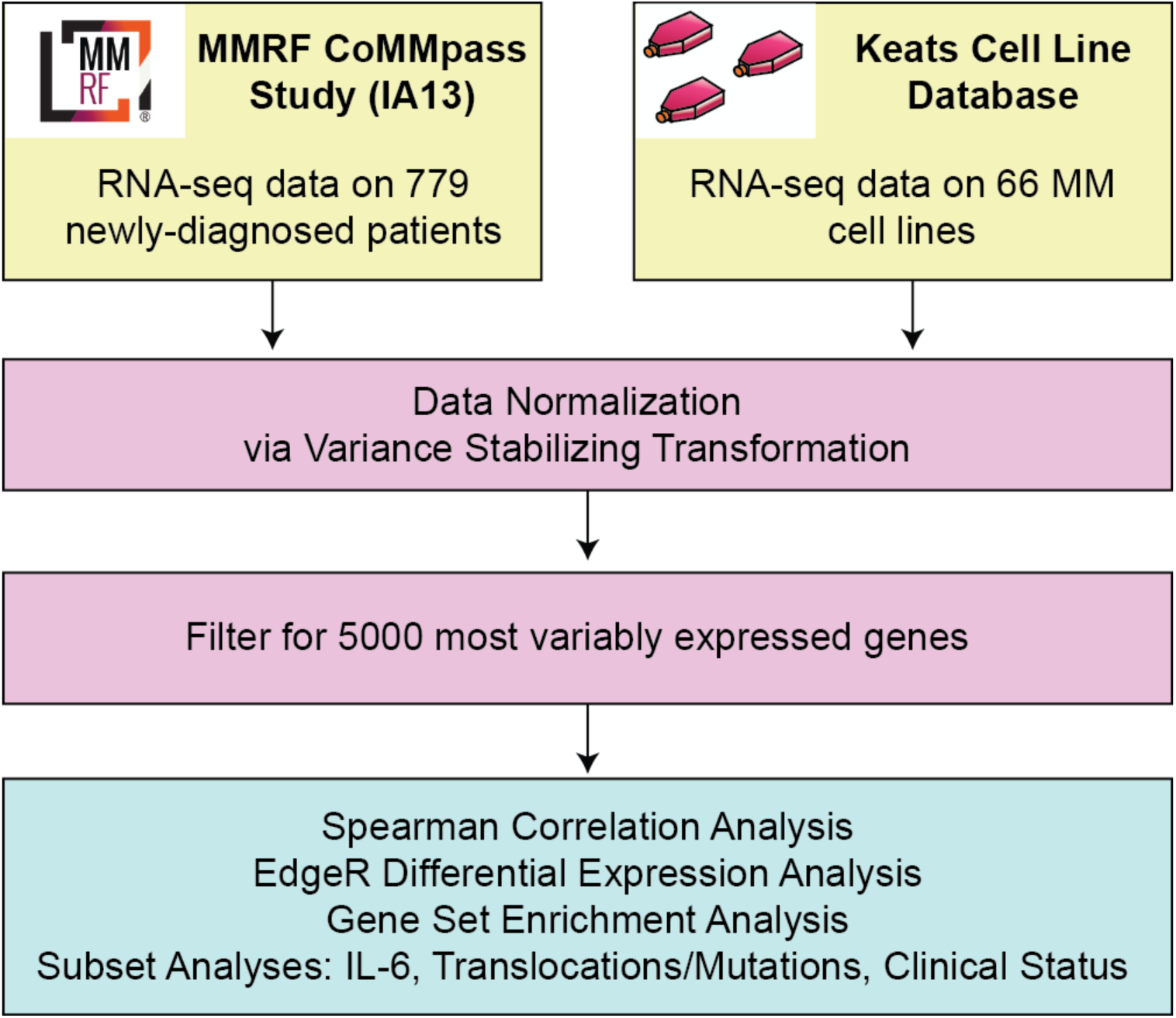
Workflow for RNA-seq based correlation analysis of multiple myeloma cell lines and patients.

**Figure 2.**
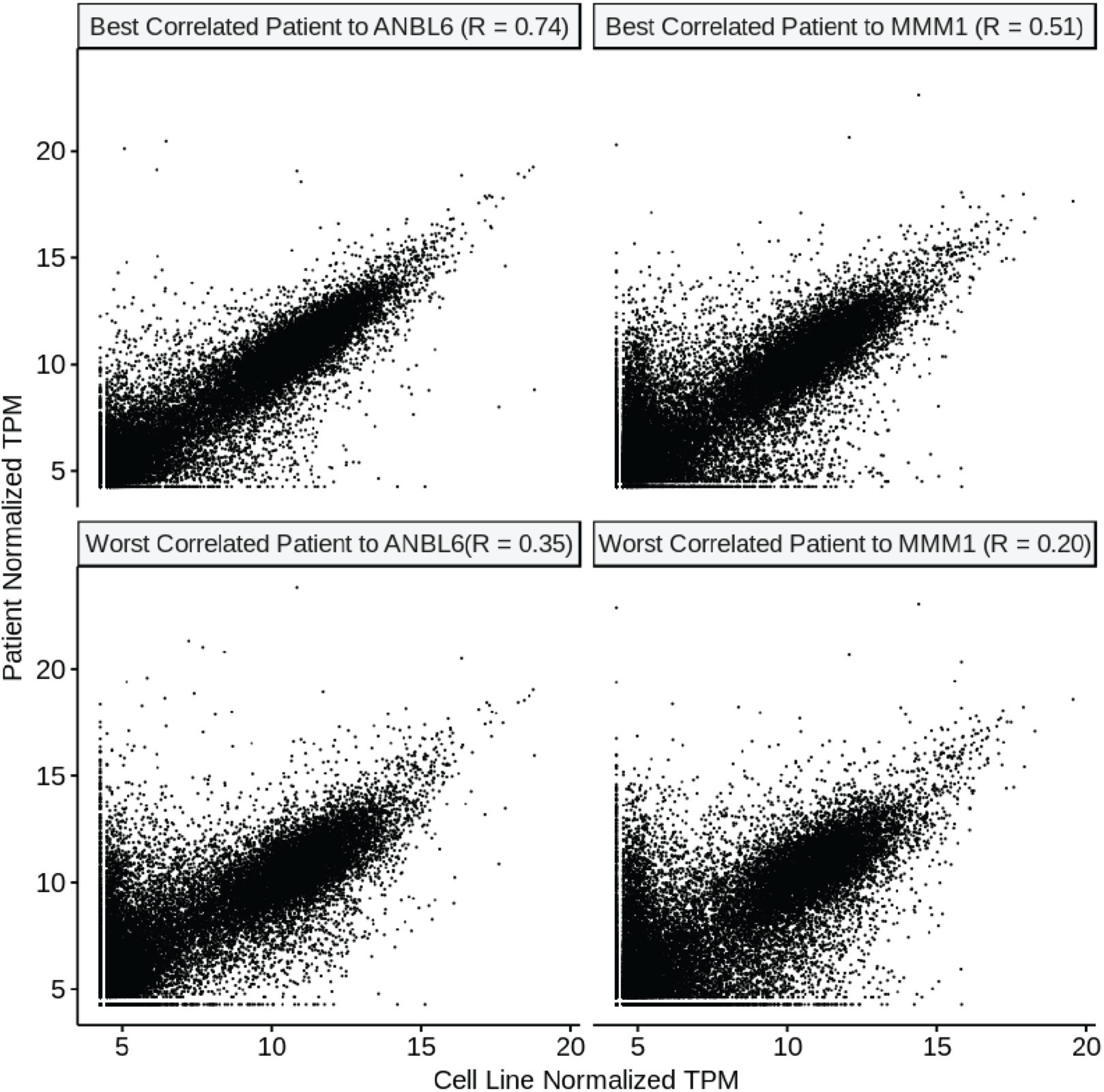
Example correlation plots for individual patients and cell lines. We show here examples of the best cell line (ANBL-6) and the worst cell line (MMM1) from our ranking in Fig. 3. Gene expression in transcripts per million (TPM) from RNA-seq data is plotted versus gene expression for the highest-correlating and lowest-correlating patient for each cell line. Similar correlations underpin the other analyses performed throughout this work.

**Figure 3.**
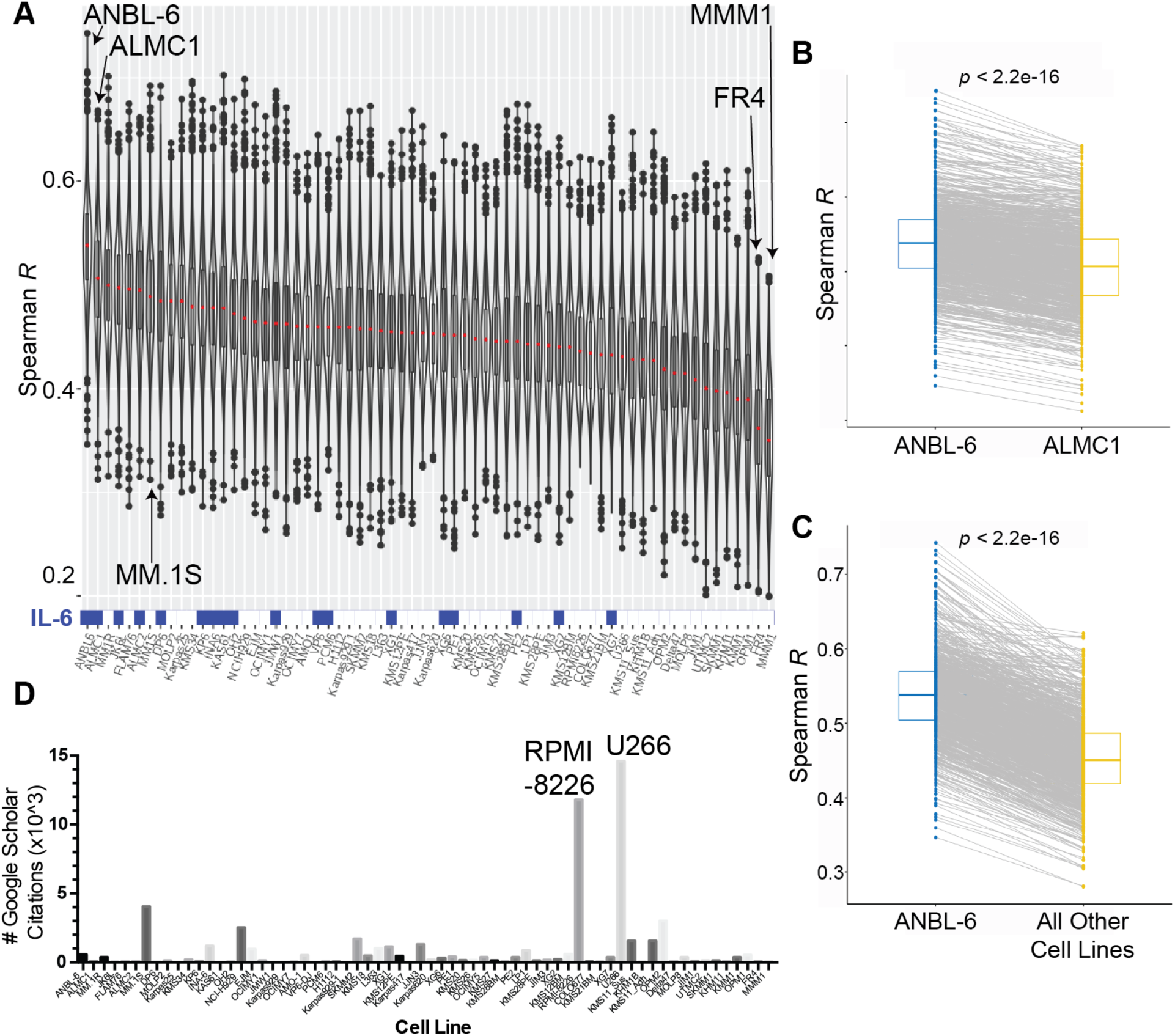
Overall MM cell line rankings reveal more and less patient-like *in vitro* disease models. **A.** Correlation analysis of the CCLE and CoMMpass data. Each sample in the violin plot corresponds to the Spearman correlation between one cell line and one primary tumor sample using the 5000 most variable genes. In the overlaid boxplot, the red center line depicts the median, the box limits depict the upper and lower quartiles, and the whiskers depict 1.5 times the interquartile range. Culture with IL-6 prior to RNA-seq analysis is indicated as blue boxes at bottom of plot. **B.** Comparison of each patient’s Spearman correlation to ANBL-6 vs. the second-ranked line, ALMC-1, demonstrates the highly significant increased correlation with ANBL-6. **C.** Similarly, essentially all patient transcriptomes correlate more strongly with ANBL-6 transcriptome than the aggregate panel of all other cell lines. All *p*-values using Wilcoxon test. **D.** The literature usage for each cell line was measured using a Google Scholar search (Oct. 2, 2019). The number of individual results from the text search of “(cell line) myeloma” is plotted per cell line and ordered per the rankings in 3A.

The violin plots in Fig. 3A are presented for each cell line in the Keats lab database, ranked by the median Spearman *R* when correlated versus each patient in the CoMMpass database. We can draw some initial conclusions from this dataset. First, it is clear that none of the MM cell lines approached a perfect representation of patient tumor, as the median *R* values range from 0.35-0.54 (i.e. far from 1). Consistent with this conclusion, Principal Component Analysis of overall transcript expression demonstrated that MM cell lines form a distinct cluster from patient tumors (Fig. S1). Second, while many of the cell lines in the middle of the ranking showed quite similar correlations to patient tumor, the cell line ANBL-6 sat atop the ranking as a notable outlier (median *R* = 0.54). In parallel, the cell lines MMM.1 and FR4 appeared markedly below other lines (median *R* = 0.35 and 0.36, respectively).

We further evaluated these findings by comparing the paired analysis of each patient tumor correlated with ANBL-6 versus the second-ranked cell line, ALMC-1 (Fig. 3B), as well as all other cell lines (Fig. 3C). In both cases, ANBL-6 led to significantly higher correlations (Wilcoxon *p* < 2.2e-16). Similarly, MMM.1 and FR4 led to significantly worse correlations versus all other lines (Fig. S2).

These results support the notion that while no MM cell line is perfect, some are still better (or worse) than others. Notably, we compared our cell line rankings to the frequency of use of MM cell lines in the literature (Google Scholar, Oct. 2, 2019) (Fig. 3D). As expected, the well-known and earliest-established^2^ lines RPMI-8226 and U-266 emerged at the top of the citation ranking. However, a quick glance revealed that these lines are not localized to the top of the patient similarity rankings. U-266, for example, despite its number 1 citation rank at approximately 14,600 mentions in the literature, actually appeared to be one of the lesser-representative lines (rank 52 of 66). RPMI-8226, with 11,800 uses in the literature, ranked 48. Our top cell line, ANBL-6, is certainly used in the literature, with 563 publications employing it, but still only comes in at number 16 in the citation rankings (**Dataset S1**). Fortunately, MMM.1 and FR4 are only rarely used in the literature (56 and 64 uses, respectively). Overall, these results indicate that frequency of appearance in the literature does not strongly predict whether a cell line actually well-represents patient tumor.

### Cell line rankings are largely consistent across laboratories

Decades of anecdotal experience have suggested that cell lines may demonstrate phenotypic “drift” when cultured in different laboratories. Recent large-scale, multi-omic studies have systematically confirmed and quantified these effects^18, 19^. Therefore, we used an orthogonal resource, the Cancer Cell Line Encyclopedia (CCLE)^20^, to evaluate whether our rankings still hold when based on RNA-seq data generated by an entirely different group. Fortunately, we were able to examine a substantial cohort of 25 overlapping cell lines between the CCLE and Keats databases (Fig. S3A).

We were encouraged to find strong consistency between the rankings generated from both cell line datasets (Fig. 4A). Upon visual inspection it is clear that the same cell lines show a notable tendency to stay near the top and bottom of both rankings. Quantitative comparison of rankings and median Spearman correlations also demonstrated high reproducibility (Fig. 4B,C). Furthermore, statistical analysis also confirmed there is no significant difference between the median correlations generated from each cell line dataset (Fig. S3B). Notably, both rankings assert that the most commonly-used lines RPMI-8226 and U-266 are relatively poor representatives of patient tumor. In contrast, among frequently-used lines, MM.1S retains one of the highest scores by both rankings. While CCLE does not include ANBL-6, the reproducibility of the overall ranking increases confidence that this top-ranked line in the Keats dataset will also show similar patient-representative gene expression patterns when used in other laboratories.

**Figure 4.**
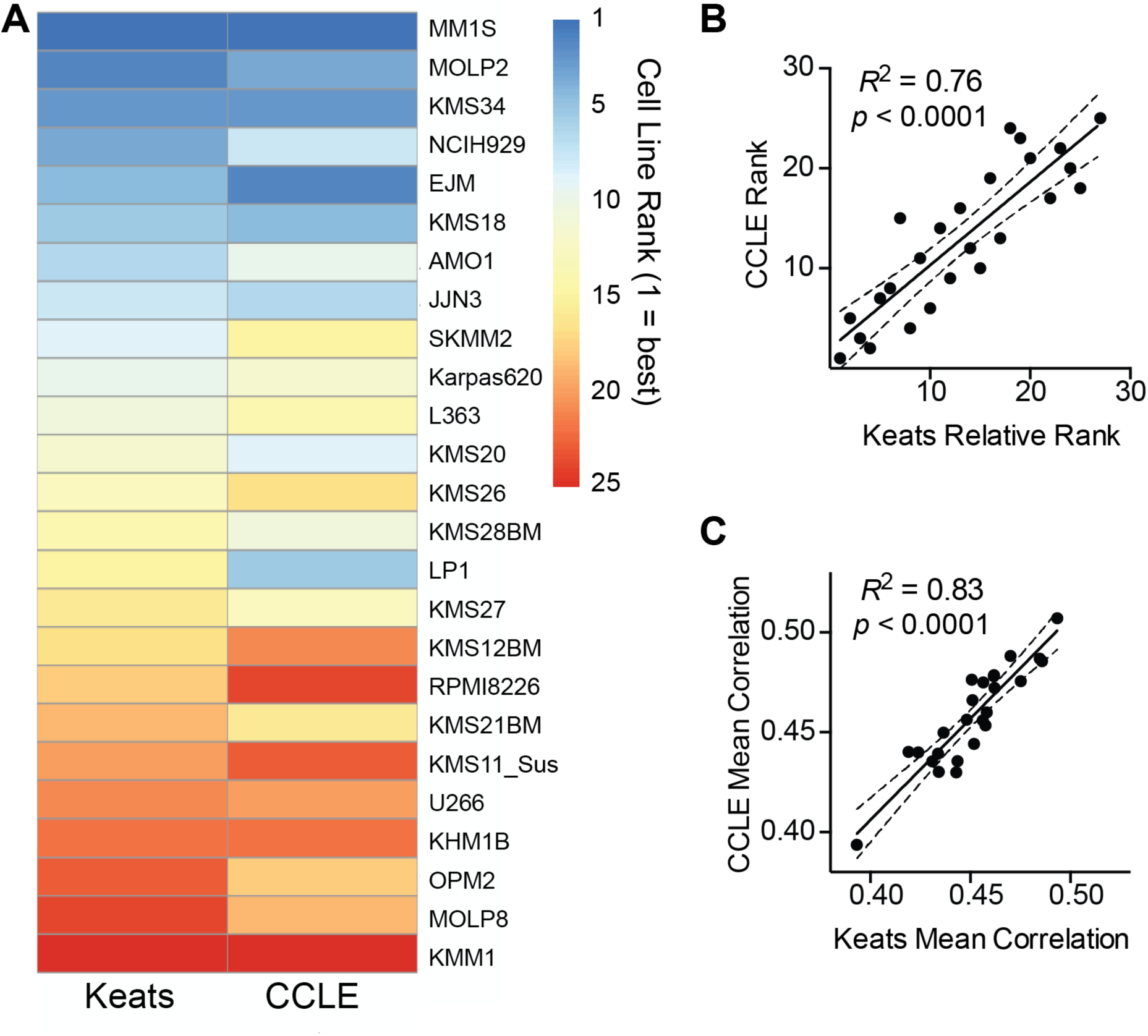
Cell line correlation rankings are largely reproducible across RNA-seq datasets from different laboratories. **A.** We performed our correlation rankings using RNA-seq data from 25 MM cell lines available in both the CCLE and Keats lab databases, with each line compared to all patients in CoMMpass. Cell lines at the top of the ranking (blue) tend to remain at the top in both rankings, and those at the bottom (red) tend to remain at the bottom in both rankings. **B.** Further supporting reproducibility, numerical rankings in CCLE and relative rankings in Keats database (numbered 1-25 to reflect rank order in overall 66 cell line ranking) are highly correlated. **C.** Similarly, mean Spearman *R* of transcriptome between each cell line and all patients, as determined from each database, is highly consistent. Linear regression displayed with 95% confidence intervals.

### Culture with IL-6 drives similarity between cell line and patient tumor transcriptome

We noted that ANBL-6, our top-ranked line, was initially characterized as being dependent on IL-6^21^. We therefore tested the hypothesis that culture of cell lines with IL-6 generally produces a more “patient-like” transcriptional signature. Indeed, we found this to be the case, where lines cultured in IL-6 in the Keats dataset showed a significant improvement in median correlation versus all patient tumors (Wilcoxon *p* = 9.5e-4) (Fig. 5A). While the absolute difference in the correlations across cell lines is modest (mean *R* = 0.48 for IL-6 cultured vs. 0.45 no IL-6), we do note that lines cultured in IL-6 show a clear enrichment in the top half of the rankings (Fig. 3A). Furthermore, while the annotated lines were cultured with IL-6 for RNA-seq analysis, many were subsequently found to not actually be dependent on this cytokine (https://www.keatslab.org/projects/mm-cell-line-characterization/cell-line-characterization-status). This finding suggests that co-culture with critical microenvironment factors can at least partially drive cell lines to a more patient-like phenotype, even if not strictly required for cell growth.

**Figure 5.**
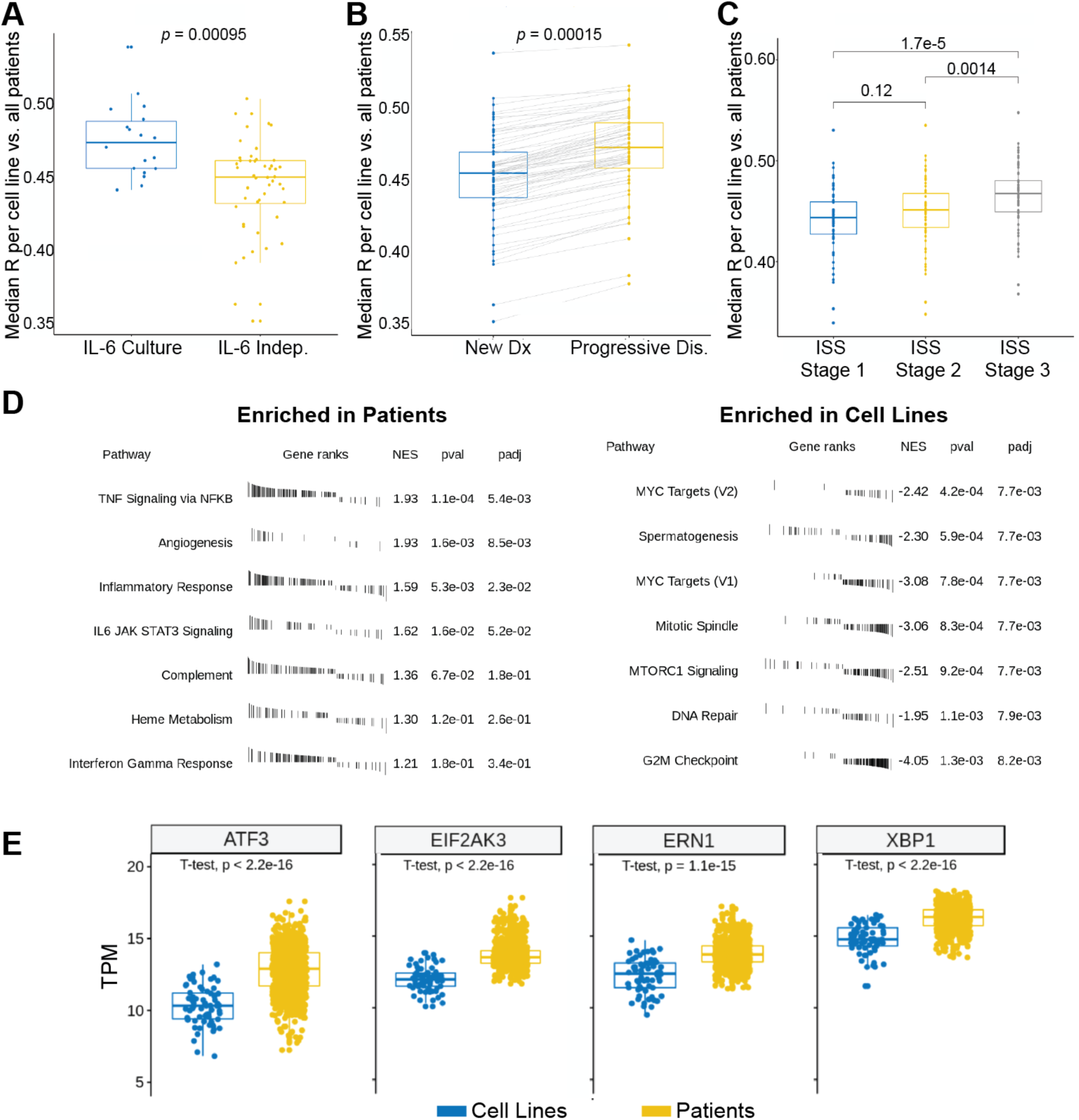
Biological factors drive increased correlations and overall differences between patient tumors and cell lines. **A.** Box plots (each dot median of a cell line compared to all patients in CoMMpass) indicates culture with IL-6 significantly increases similarity of cell lines to tumors. **B.** The cohort of CoMMpass patients with progressive disease showed increased similarity to cell lines versus newly-diagnosed. **C.** Increased International Staging System (ISS) grade at diagnosis leads to more similarity to cell lines. **D.** Gene Set Enrichment Analysis (GSEA) reveals immune signaling signatures significantly enriched in patient tumors, whereas signatures of proliferation and oncogenesis are enriched in cell lines. Performed using cutoff of differentially-expressed genes at Log2 fold-change >|1|, False Discovery Rate < 0.01. **E.** Comparative expression of selected genes (in transcripts per million, TPM) related to unfolded protein stress identifies differences in cell lines and patients. *p*-values by Wilcoxon test.

### Poor-prognosis clinical features drive similarity between cell lines and patient tumors

We next surmised that MM plasma cells able to grow *in vitro* are likely selected for increased proliferative capacity. As a corollary, patients with more aggressive disease may therefore have tumors with more similarity to cell lines. To test this, we first analyzed newly-diagnosed patients in CoMMpass based on International Staging System (ISS) stage at diagnosis (Fig. 5B). We indeed found that the poorest-prognosis patients, at ISS stage 3, had tumors with transcriptional profiles significantly more similar to all cell lines together (*p* = 0.0014 and 1.7e-5 vs. stage 2 or stage 1, respectively). Similarly, patients with higher M-spike at baseline also showed greater tumor similarity to cell lines (Fig. S4).

We also evaluated the smaller cohort of patients with progressive disease in CoMMpass (*n* = 81) and obtained their overall transcriptional correlations to all cell lines, in comparison to our prior newly-diagnosed analysis (*n* = 779). Here we also found significantly increased correlation with the relapsed versus newly-diagnosed patients (*p* = 0.0015) (Fig. 5C). Taken together, our results provide quantitative support for the notion that MM cell lines more closely resemble more aggressive, poor-prognosis disease states rather than the “typical” newly-diagnosed myeloma patient.

### Specific biological signatures differentiate cell lines and patient tumors

While these results help clarify the basis of MM cell line phenotypes, we next used Gene Set Enrichment Analysis (GSEA) to further delineate biological signatures that most distinguish cell lines and patient tumors (Fig. 5D). We found that signatures relating to cell cycle/proliferation, mTOR signaling, and MYC targeting were significantly upregulated in cell lines versus patient samples. These findings underscore how MM cell lines have adapted to the setting of rapid, cell-autonomous proliferation *in vitro*. In contrast, patient tumors showed increased signatures of immune and microenvironment signaling including IL-6/JAK/STAT3 signaling, interferon response, TNF signaling, and complement. Similar findings were obtained with Gene Ontology (GO) analysis (Fig. S5). These results illustrate the importance of MM immune microenvironment effects in driving human *in vivo* transcriptional phenotypes. These results also suggest that exposing cell lines to more of these microenvironmental factors, including, but not limited to, IL-6, may assist in further driving a patient-like signature *in vitro*.

Furthermore, it has been reported that MM cell lines frequently downregulate immunoglobulin synthesis compared to patient tumors^22, 23^. We also observed this phenomenon comparing expression of all transcripts of patients vs. cell lines (Fig. S6 and **Dataset S3**). We hypothesized this decreased protein load would lead to a decrease in baseline unfolded protein stress in cell lines. Indeed, we also found decreased cell line expression of genes that govern protein homeostasis and unfolded protein stress in the endoplasmic reticulum, such as ATF3, EIF2AK3/PERK, XBP1, and ERN1/IRE1 when compared to patients (Fig. 5E). Given the prominent role of misfolded immunoglobulin burden in proteasome inhibitor-induced apoptosis^24–26^, this result may partially explain why MM cell lines are not markedly more sensitive to bortezomib than many solid tumor cell lines, whereas among cancer patients only those with MM have shown strong clinical responses to proteasome inhibition^27^.

### MM cell lines carry unique mutational signatures compared to patient tumor

Over the past 10 years, large-scale whole genome and whole exome sequencing studies have revealed numerous mutations found recurrently in MM^28–30^. These findings follow prior cytogenetic studies which have found large-scale chromosomal aberrations, including both translocations and copy-number variants, that drive differential patient prognosis and are routinely tested in the clinical setting^31^.

Here we took advantage of whole exome sequencing data in CoMMpass and the Keats lab cell line database to investigate the relative frequency of mutations in both sample sets (Fig. 6A). We first note that activating mutations in the most recurrently-altered oncogenes in patients, *KRAS* and *NRAS*, are mutated at similar frequency in both cell lines and patient samples (30.9% vs. 25.1% *KRAS*, 20.1% vs. 21.1% *NRAS*, respectively) demonstrating consistency between these key sequence variants. *TP53* mutations were markedly more common in cell lines (55.9% vs. 4.1% in patients), potentially consistent with the more aggressive growth phenotype of cells *in vitro*. Other commonly-mutated genes in patient tumors, as characterized by Walker et al.^30^, generally show similar mutation frequencies in cell lines and patients (Fig. S7). However, beyond these well-known genes, we also noted several unexpected genes that were mutated at high frequency in cell lines but infrequently in patients (Fig. 6A and **Dataset S2**). These included *OLFM1*, *MUC16*, *MUC3A*, *MUC6*, *MUC4*, *IGSF3* and *ZNF527*.

**Figure 6.**
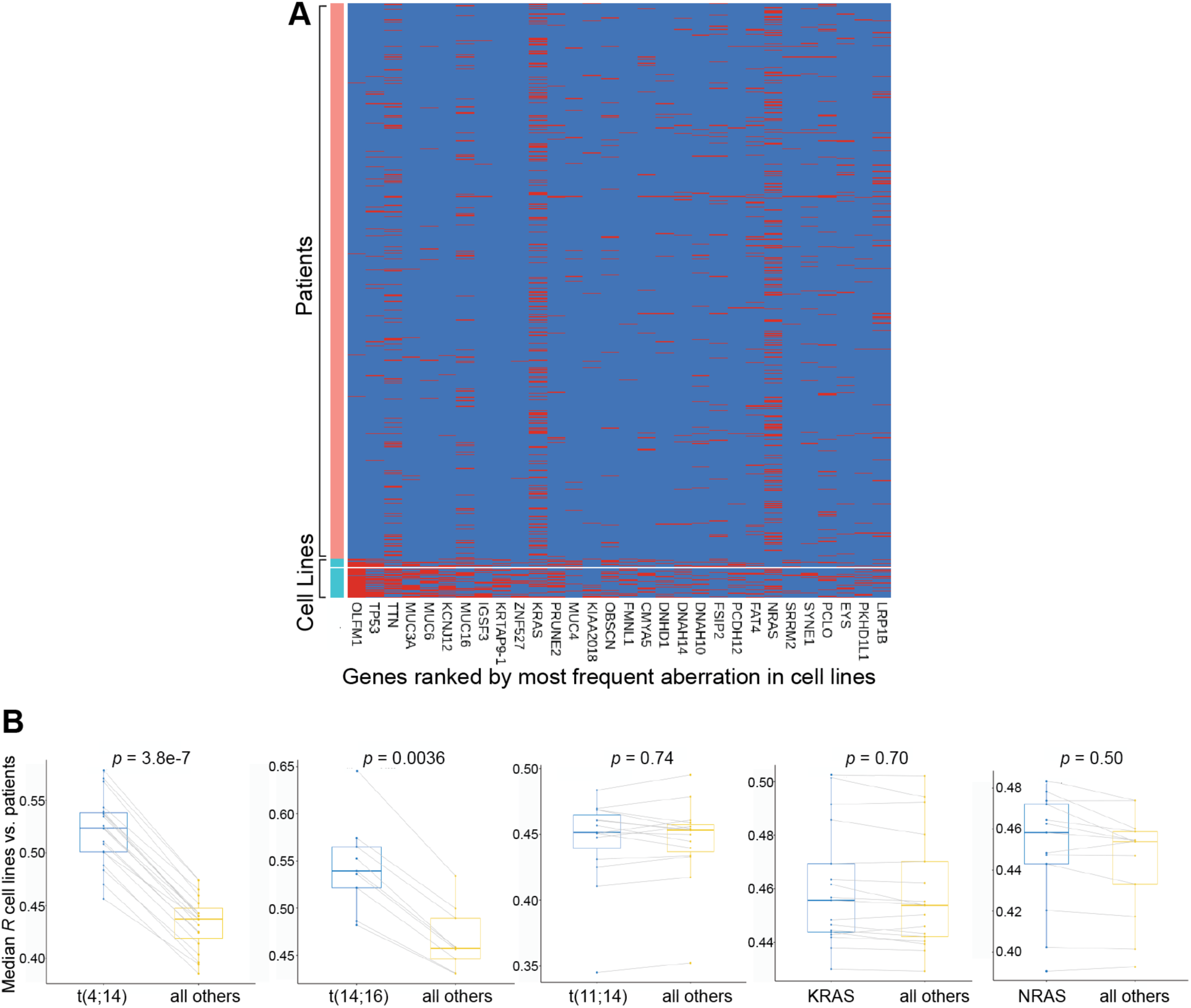
Integrating genomic with transcriptomic analysis to compare patient tumor vs. cell lines. **A.** Analysis of exome sequencing data in the Keats cell line database and patient tumor in CoMMpass reveals genes frequently mutated in cell lines with limited, if any, identification of related variants in patient samples. Red = mutation detected compared to hg19 reference genome. Blue = no mutation. **B.** Subset analysis of correlation profiling when matching canonical myeloma genomic lesions (three *IGH* translocations, *KRAS*/*NRAS* mutations). Each dot reflects the median Spearman correlation of each cell line carrying the specified genomic lesion correlated versus CoMMpass patients with or without the noted genomic lesion. Box plot shows median and interquartile range. *p*-values by Wilcoxon test.

The biological significance of these mutations in driving MM cell-line specific phenotypes is not immediately clear. Initial analysis of expression data suggest most of the genes are expressed at very low levels, with the exception of *ZNF527* (Fig. S8). *MUC16* is a relatively large gene (total gene length 133 kb per hg19 reference genome) with missense mutations throughout the sequence (**Dataset S2**). It is unclear if these alterations relate to tumor biology or if they are randomly found due to the large size of this gene and general genomic instability of cancer cell lines. This finding would be similar to *TTN* (281 kb), encoding the giant muscle protein titin, where mutations are frequently observed in other cancer genomic studies and are not thought to carry clinical relevance^32^. In contrast, specific recurrent mutations are found frequently in the other genes noted above (**Dataset S2**) but very rarely if at all in patients. This finding raises the hypothesis that these variants are indeed selected for in the context of promoting *in vitro* growth. In particular, the enrichment in MUC family glycoprotein genes in this small gene list is quite striking. The related gene *MUC1*, while not included in the list here, has been proposed as a potential oncogene in MM^33^. Therefore, we cannot exclude the possibility that these recurrent mutations in MUC family genes indeed carry some role in promoting autonomous plasma cell growth, even at low gene expression levels.

### Matching of common MM genomic aberrations does not always lead to increased cell line-patient transcriptional similarity

In MM research it is common to use cell lines with particular genomic lesions as proxies for biological features for patients with the same aberrations. We next tested whether some of these most-common genomic aberrations - translocations (11;14), (4;14), and (14;16), as well activating mutations of *NRAS* and *KRAS* (codons 12/13/61) - improved global transcriptomic correlations when matched between cell lines with patients carrying the same lesion. Our analysis confirmed that matching of t(4;14) and t(14;16) cell lines indeed improved correlations to patients with the same alteration as compared to those without (*p* = 3.8e-7 and 0.0036, respectively) (Fig. 6B**, left**). However, we saw no significant increase when matching t(11;14) or activating *RAS* mutations (Fig. 6B**, right)**. While these results by no means refute the utility of extrapolating findings from cell lines with specific aberrations to patients with the same genotype, they do surprisingly indicate that these latter genotypes do not lead to broad-scale increases in the global cellular transcriptome similarity based on presence of the same lesion.

### ANBL-6 is appropriate for disseminated in vivo MM modeling

Our overall rankings (Fig. 3A) suggest that ANBL-6 should be incorporated more frequently into MM studies. Toward more widespread use of ANBL-6, one potential drawback for MM *in vitro* studies is the cost of recombinant IL-6. We therefore titrated IL-6 and found that over 72 hours, a minimal concentration of 0.1 ng/mL was able to support equivalent proliferation to 100 ng/mL (Fig. S9), consistent with earlier results^34^. No proliferation was observed in the absence of IL-6, confirming IL-6 dependence.

Furthermore, while ANBL-6 has been used in many prior studies (Fig. 2D), the vast majority of the efforts were purely *in vitro*. Disseminated orthotopic xenograft models of MM, where luciferase-labeled plasma cells are intravenously (I.V.) implanted in NOD *scid* gamma (NSG) mice, may carry significant advantages for preclinical modeling if tumor cells home to hematopoietic tissues including bone marrow^35^. In this context, cells will proliferate and respond to therapy in a microenvironment more akin to that present in patients.

To our knowledge, it has not been tested whether ANBL-6 homes to hematopoietic tissues in a disseminated mouse model after I.V. implant. We therefore used lentiviral transduction to stably express luciferase in ANBL-6 cells and injected 1e6 cells into a pilot cohort of four NSG mice. In parallel, we injected 1e6 luciferase-labeled MM.1S cells into a separate cohort as a control, as these cells are well-known to home to bone marrow in NSG mice^36^. Encouragingly, we found that ANBL-6 showed an identical pattern of distribution as MM.1S, with implantation primarily to the spine, sternum, and hindlimbs (Fig. 7A). However, *in vivo* growth kinetics and overall murine survival were significantly prolonged compared to MM.1S (Fig. 7B,C). Therefore, ANBL-6 may perhaps serve as a valuable *in vivo* model for a more indolent form of MM, rather than highly-aggressive disease as represented by most disseminated MM cell line models.

**Figure 7.**
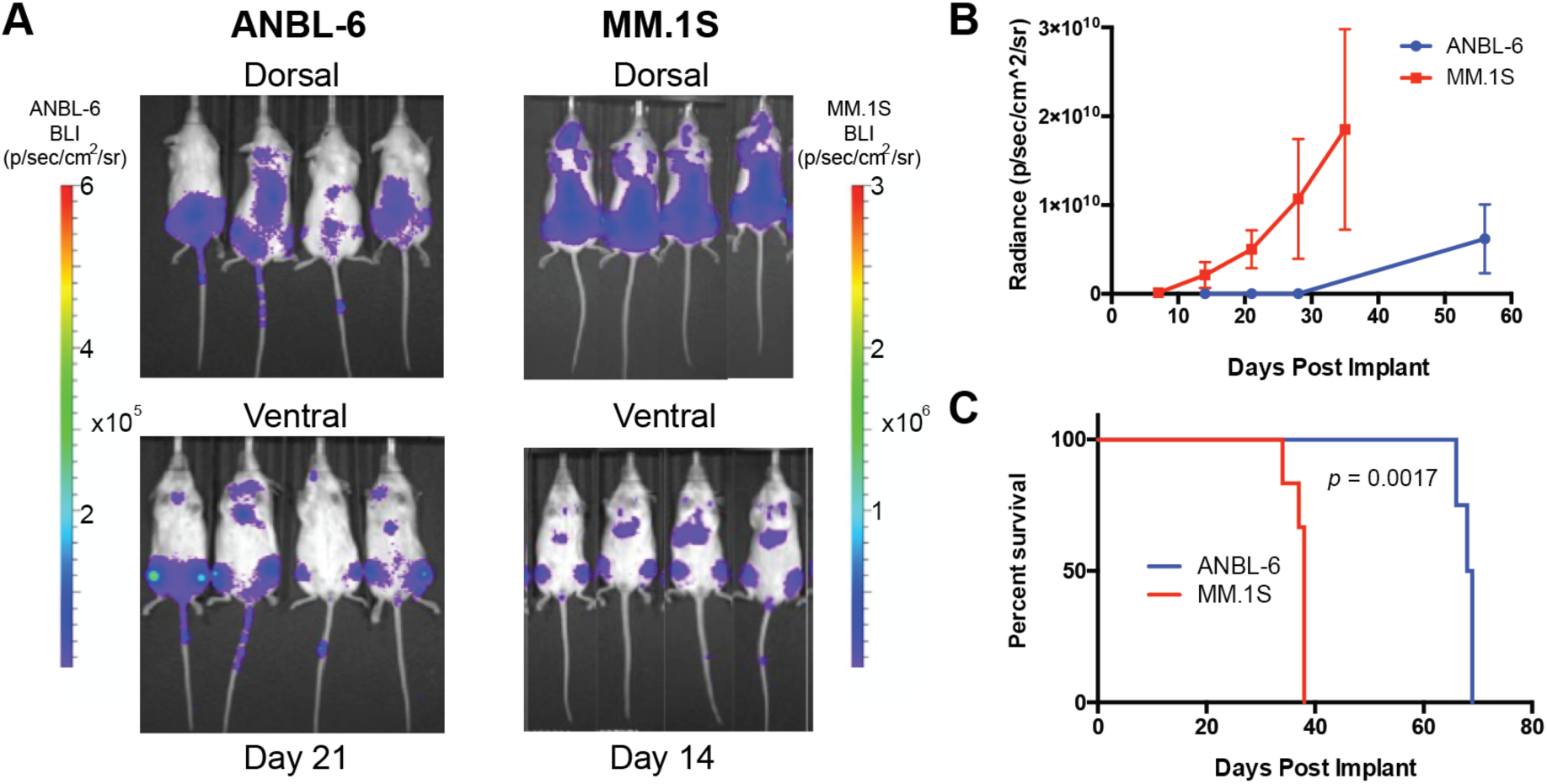
Investigating the potential of ANBL-6 for preclinical *in vivo* modeling in myeloma. **A.** 1e6 luciferase-labeled ANBL-6 and MM.1S cells were implanted via tail vein injection into NSG on the same date. Bioluminscence imaging (BLI) data is shown at noted dates post-implant. Both cell lines show identical localization, primarily to long bones of hindlimbs and to the spine. Note scale bar with lower intensities for BLI signal in ANBL-6. **B.** Quantification of BLI signal (*n* = 4 in ANBL-6 arm; *n* = 6 in MM.1S arm). **C.** Survival of MM.1S-implanted mice vs. ANBL-6 implanted mice illustrates more indolent course of ANBL-6 *in vivo*. *p*-value by Log-Rank test.

## Discussion

Here we present the first large-scale quantitative comparison of MM cell lines and primary patient tumors. Our results, using global transcriptome profiles as a proxy for overall biological state, quantitatively confirm long-standing suspicions that MM cell lines are indeed very different than patient tumors. Through these analyses, we describe biological factors that drive increased or decreased similarity between cell lines and patients, as well as outline strategies to potentially improve the quality of *in vitro* studies toward representing *in vivo* patient disease.

While our results reveal that all MM cell lines are biologically divergent from patient tumors, we argue that it is still certainly possible to improve the quality and relevance of *in vitro* studies by incorporating our rankings and other findings here. Even though many of our comparative analyses (IL-6 co-culture, relationship to progressive disease, etc.) show relatively modest absolute increases in global transcriptional correlation, these increases remain highly statistically significant and involve biologically relevant changes over hundreds of genes.

We note that our analysis here is only enabled by recent, large scale RNA sequencing-based studies in MM. Prior microarray-based expression profiling analyses of patient samples^37^, for example, do not readily allow for similar robust normalization and quantification approaches when comparing across different datasets. In parallel, though, our findings here are of course limited by the fact that they are solely based on the transcriptome. Other ‘omic signatures (metabolomics, proteomics, etc.), or, alternatively, curated individual markers, may further refine and extend these results. However, given current technologies, we would argue that transcriptional profiles are the best way to directly assess the relationship of these cell line models to patient disease.

Similar to our prior pan-cancer analysis of the TCGA and CCLE^13^, here we identify MM cell line models that appear both more and less representative of patient disease. In particular, our results suggest that the two most widely-used MM cell lines in the literature, U-266 and RPMI-8226, are actually some of the least-representative of patient disease (Fig. 3). In contrast, we find that the third-most-used cell line, MM.1S, does appear to be one of the better models available. Importantly, these findings were reproducible across two different datasets (Fig. 4).

Our results specifically indicate that the cell line ANBL-6 sits significantly above all other cell lines in terms of patient similarity. ANBL-6 was isolated from peripheral blood of a relapsed MM patient and initially characterized as having typical malignant plasma cell immunophenotype, a (14;16) translocation, and lambda light chain secretion^21^, and later shown to have wild-type *NRAS* and *KRAS* sequences^10, 34^. We confirmed that ANBL-6 showed consistent proliferation even at a minimal, cost-effective IL-6 concentration. Notably, our prior results suggest that other factors within the NSG murine marrow microenvironment may be able to partially compensate for the lack of cross-talk between murine IL-6 and human IL-6 receptor^36^, thereby allowing for *in vivo* proliferation of ANBL-6. Furthermore, we found that co-culture with IL-6 appears to drive more patient-like phenotypes across all cell lines (Fig. 5A), and IL-6-mediated signaling is a prominent transcriptome signature enriched in patient tumors (Fig. 5D). We therefore propose two readily-implemented actions based on our results: 1) more widespread use of ANBL-6 with decreased use of RPMI-8226 and U-266; 2) more common use of IL-6 in culture media, potentially even for lines not strictly dependent on this cytokine.

The overall conclusions here will not necessarily be surprising to MM researchers given years of anecdotal experience and knowledge of MM biology. However, like recent studies systematically investigating differences in cell line phenotype across different laboratories^18, 19^, these results are important to provide quantifiable metrics to compare cell lines and patient tumors, and potentially provide benchmarks for the development of new lines. The analyses here stand as a resource with widespread utility to the MM community and lead to specific recommendations for alterations in research practice.

## Acknowledgments

We thank Drs. Brian Van Ness, Jonathan Keats, Tom Martin, Nina Shah, Sandy Wong, and Jeffrey Wolf for insightful discussions. We thank the Multiple Myeloma Research Foundation for sponsoring the cell line transcriptome and CoMMpass studies. We thank Dr. Benjamin Barwick for providing analysis of translocation status in patient samples. Funding for this study was provided by the UCSF Stephen and Nancy Grand Multiple Myeloma Translational Initiative, NIH DP2 OD022552, K08 CA184116, and R01 CA226851 (to A.P.W.), K01 LM012381 (to M.S.), and UCSF Helen Diller Family Comprehensive Cancer Center P30 CA082103 (to B.C.H.).

## Author Contributions

V.S., K.Y., M.S., and A.P.W. conceived and designed the study. V.S. and K.Y. performed bioinformatics analyses. I.D.F., O.G., M.A.N., and B.C.H. developed luciferase-labeled ANBL-6 cell line and performed ANBL-6 murine experiments. V.S., K.Y., M.S., and A.P.W. wrote the manuscript with input from all authors.

## Supplementary Materials

### Supplementary Methods

#### Data collection and normalization

The cell line gene expression file (HMCL66_HTSeq_GENE_Counts.txt) was downloaded from the Keats Lab repository (https://www.keatslab.org/data-repository). For patients the gene expression file (MMRF_CoMMpass_IA13a_E74GTF_HtSeq_Gene_Counts.txt gene) was downloaded from the CoMMpass study (https://research.themmrf.org/). Information on how gene expression data for patients was aligned and count based expression estimates calculated can be found in the (MMRF_CoMMpass_IA13_Methods.pdf) file. We used the 48637 Ensembl IDs that were in both the Keats dataset and the CoMMpass dataset for our analysis.

The file (HMCL69_Preliminary_Mutations_Samtools.xlsx) was utilized to determine which cell lines were mutated for mutational subtype analysis as well as for mutational frequency and the file (MMRF_CoMMpass_IA13a_IGV_All_Canonical_Variants.mut) was used to gather patient data on the same. Translocations of cell lines were determined from the (Myeloma_Cell_Line_Characteristics.csv) file downloaded from (https://www.keatslab.org/myeloma-cell-lines/hmcl-characteristics). For patients this data was derived from the (MMRF_CoMMpass_IA13a_Delly_Structural_Calls.txt) file (see below). Patient annotations for progressive disease and Serum M-protein levels came from the (MMRF_CoMMpass_IA13_PER_PATIENT_VISIT.csv) file. Patient annotations for ISS staging came from the (MMRF_CoMMpass_IA13_PER_PATIENT.csv) file.

Data was normalized using vst, the DESeq2 (Love *et al.*, *Genome Biol* (2014) 15:550) wrapper for varianceStabilizingTransformation function in R.

#### Correlation Analysis

We filtered low count genes by only retaining genes that had greater than 1 counts per million (CPM) in 2 or more samples, leading to analysis of almost exclusively protein-coding genes. We then normalized by variance stabilizing transformation before utilizing the 5000 most variably expressed genes as determined by the interquartile range (IQR) to compare cell lines and newly diagnosed patient samples using Spearman’s rank correlation.

We chose the 5000 most variable genes based on previous studies (Yu et al., *Nat Commun* (2019) 10:3574; Chen et al., *BMC Med Genom* (2015) 8:S5). However, we did also analyze the effect of instead choosing the top 10000 most variably expressed genes. In this wider analysis we found the correlation coefficients generally increased but there was an insignificant shift of overall rankings vs. the top 5000 (not shown). For consistency with prior studies in the field we therefore chose to complete our analyses using the 5000 most variable genes.

#### Subtype Analysis

Translocation and mutation information for cell lines was annotated by the Keats Lab. Mutational information for patients were also found annotated in the CoMMpass dataset. For patient translocation information, we utilized the DELLY VCF files to determine translocations as previously described (Barwick *et al.*, *Nat Commun* (2019) 10:1911). Translocations with homology of 80% or more in any 100bp window within 1 kb of the translocation breakpoint were removed. Average mappability (determined from ENCODE30 20bp mappability tracks) of less than 20% across 1 kb on either side of the translocation breakpoint were removed. Finally, translocations were visually inspected and compared to matched normal tissue and translocations with sequencing anomalies were removed.

ISS staging, serum M protein levels, and progressive disease annotation were also annotated in the CoMMpass database.

For subsets of patients based on ISS staging, serum M-protein levels, and those annotated as having progressive disease in CoMMpass, we compared mean correlation coefficients of patients in each clinical category to all cell lines. For translocation and mutational analysis we correlated cell lines annotated as carrying each genomic aberration, per Keats lab data, to patients annotated as either having or not having the same aberration. The Wilcoxon rank-sum test was used to assess the differences between the groups.

#### CCLE Cross-Check

Correlation analysis was done for overlapping 25 cell lines utilizing the same 5000 genes as determined to be most variably expressed. The cell line gene expression file (CCLE_RNAseq_genes_counts_20180929.gct) was downloaded from the CCLE data repository (https://portals.broadinstitute.org/ccle/data). We conducted a Spearman’s rank correlation for all newly diagnosed patients utilizing the count-based expression estimates as derived from CCLE data and then compared the rankings and mean correlations of the cell lines as analyzed with the original Keats lab data.

#### Gene Set Enrichment and Gene Ontology Analysis

Differentially expressed genes were determined by the likelihood ratio test method with upper quartile normalization as outlined in the edgeR user’s guide. We ranked our genes by log-fold change and conducted our analysis on the 50 hallmark gene sets available on MSigDB (http://software.broadinstitute.org/gsea/msigdb/index.jsp) utilizing the fgsea R package. We considered a gene set to be up or down regulated using log2-fold change > |1| at False Discovery Rate < 0.01. For GO analysis we used the clusterProfiler package and only considered differentially expressed genes with adjusted *p*-value less than 0.05. We performed an overall analysis for patients and cell lines considering all GO annotations (molecular function, biological process, and cellular component) with redundant GO terms being removed using the simplify function in the clusterProfiler package.

#### Exome Sequencing Analysis

As annotated in CoMMpass IA13 processed and variant-called exome data, for patients we filtered for all mutations that were annotated as being a missense variant, frameshift variant, stop gained, stop lost, start lost, disruptive in-frame insertions, and deletions so as to narrow our analysis to only deleterious mutations. For cell lines we similarly utilized annotated mutations as annotated in Keatslab processed exome data. Then we filtered for genes which we had information in both cell lines and patients. We present separately the 30 most frequently mutated genes in cell lines as well as genes annotated by Walker et al. (*Blood* (2018) 132:537) as potential driver mutations in myeloma.

#### Literature results using Google Scholar search

For each cell line in the Keats lab dataset, a Google Scholar (scholar.google.com) search was performed on Oct. 2, 2019 using the term “[cell line] myeloma”. The number of unique results returned for this search is reported in Fig. 3D. For cell lines with two versions derived from different tissues from the same patient (for example, KMS-28PE and KMS-28BM) the search only used the common name of the cell line (“KMS28”) and the same result number is plotted for both cell line versions.

#### Cell Culture Conditions

ANBL-6 cells were a kind gift of Dr. Brian Van Ness at the University of Minnesota. Identity was verified by DNA genotyping and confirmed as mycoplasma-free. ANBL-6 were maintained in complete media with RPMI-1640 (RPMI-1640; Gibco) supplemented with 10% fetal bone serum (FBS; Gemini), 1% penicillin-streptomycin University of California San Francisco (UCSF), 2 mM L-Glutamine (UCSF), and 2 ng/mL IL-6 (ProSpec) with 5% CO2.

#### Generation of luciferase-labeled ANBL-6

Cell lines stably expressing enhanced firefly luciferase (effLuc) to enable *in vivo* bioluminescence imaging were generated using standard lentivirus transduction methods. Briefly, lentivirus was produced using HEK293T cells transfected with a mixture of transfer plasmid (encoding an effLuc, mCherry, Neomycin-resistance gene expression cassette) and second-generation lentiviral packaging plasmids while cells were ∼80% confluent. Transfection of lentivirus plasmids was performed using polyethethylenimine (Transporter-5, Polysciences, 26008-5) at a 4:1, PEI:DNA mass ratio. Virus containing supernatant was harvested after 72-hours and filtered using a 0.45uM syringe filter, then stored at −80°C until used. Transduction of cells was performed by the addition of 1ml of viral supernatant to 1-2E6 cells, supplemented with 8ug/ml polybrene (EMD Millipore, TR-1003-G), then spinfecting at 1000 RCF for 2 hours at 33°C. One day after spinfection the viral supernatant was replaced with fresh media and cells were allowed to recover for several days to allow for transgene expression prior to FACS sorting to achieve a uniform population of effLuc expressing cells. Construct encoding effLuc and mCherry was a kind gift of Dr. Diego Acosta-Alvear, UCSF.

#### IL-6 titration studies

1e3 ANBL-6 cells were seeded per well in 384 well plates (Corning) using the Multidrop Combi (Thermo Fisher) and incubated for 72 hr with the indicated concentration of IL-6 (ProSpec) in quadruplicate. CellTiterGlo (Promega) was used to measure viable cells per well at 72h and compared to baseline analysis at 0h prior to IL-6 treatment, with plate imaging on the Promega GloMax.

#### ANBL-6 murine studies

1e6 ANBL-6-luc or MM.1S-luc cells, stably expressing luciferase, were transplanted via tail vein injection into 4 and 6 female NOD.Cg-*Prkdc^scid^ Il2rg^tm1Wjl^*/SzJ (NSG) mice, respectively. 6-8 weeks old NSG mice were obtained from in-house breeding stocks at UCSF Preclinical Therapeutics Core (PTC) Facility. Tumor burden was assessed through bioluminescent imaging in the UCSF PTC on a Xenogen In Vivo Imaging System (IVIS) at the time points indicated in Fig. 7B. Survival is denoted by the time to development of symptomatic myeloma, at which point sacrifice is required per animal welfare guidelines. All studies were approved by the UCSF Institutional Animal Care and Usage Committee.

## Supplementary Figures

**Figure S1.**
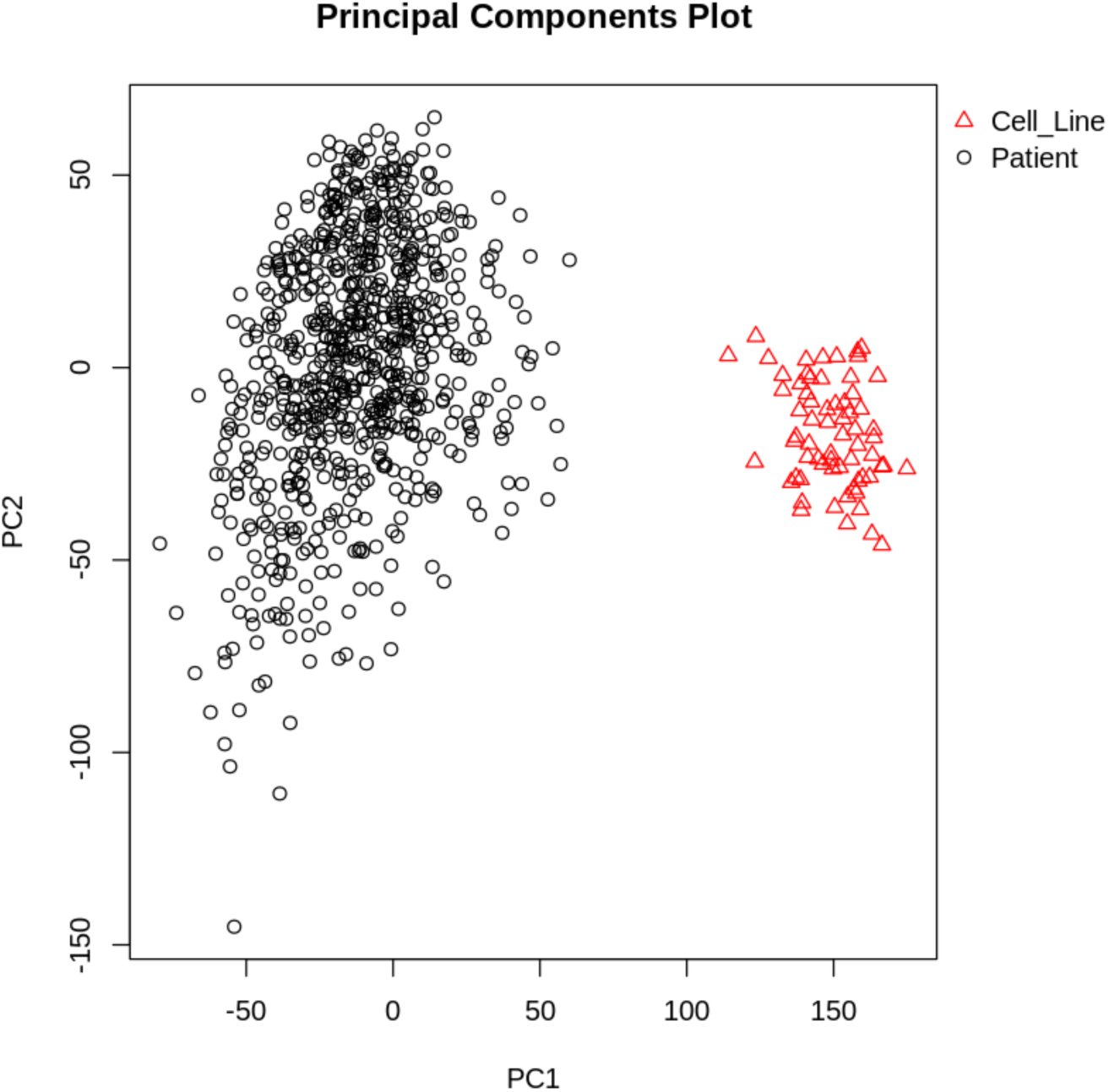
Principal Component Analysis illustrates separation between patient and tumor transcriptomes. Performed based on 5000 most variable genes after variance stabilizing transformation.

**Figure S2.**
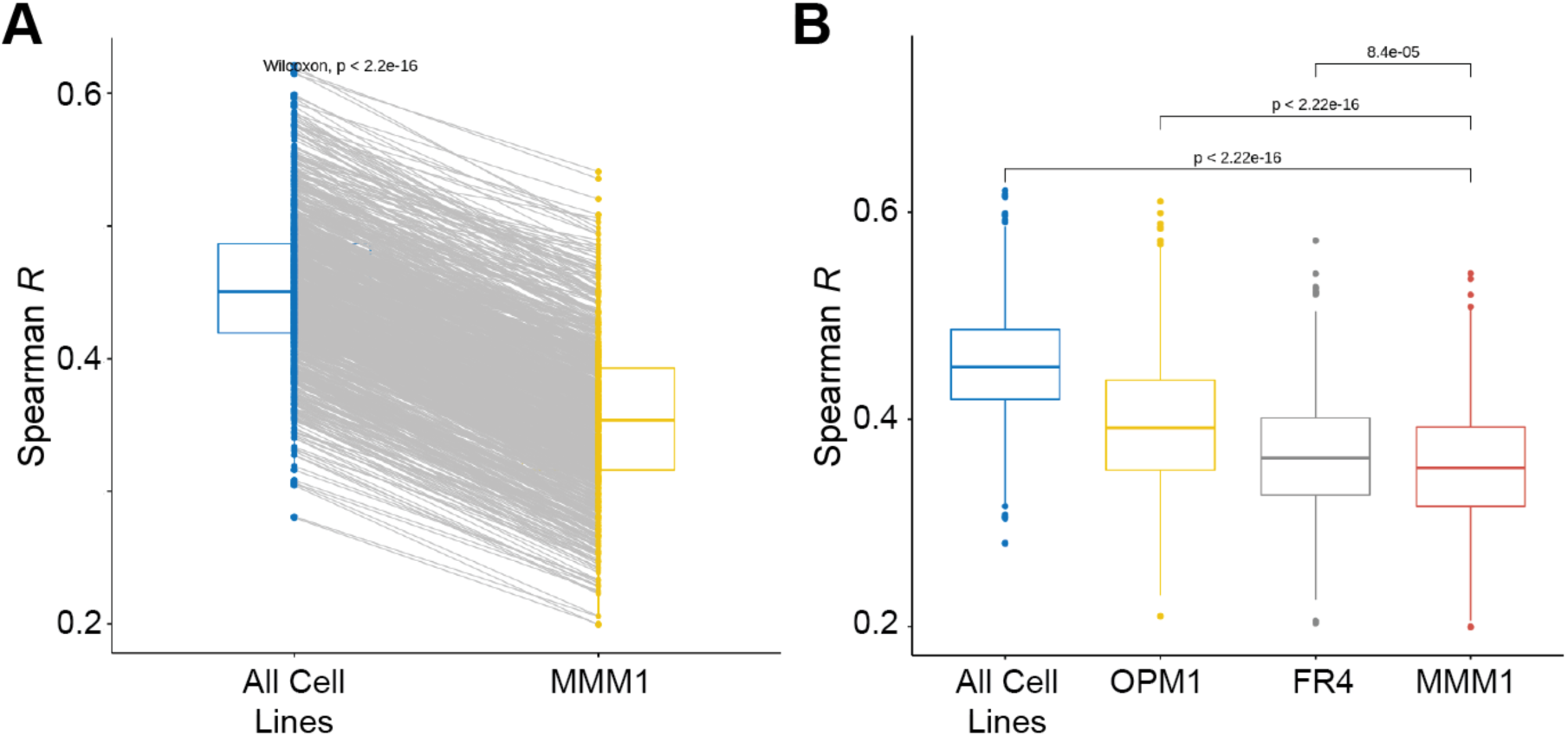
MMM1 is the lowest-ranking cell line. **A.** Analysis across all patient correlations versus this lowest-ranking line demonstrates significantly decreased patient representation of MMM1. **B.** Comparative analysis of the bottom three cell lines (OPM-1, FR4, MMM1) demonstrates that FR4 and MMM1 are significantly less representative of patient tumor than even OPM1. *p*-values by Wilcoxon test.

**Figure S3.**
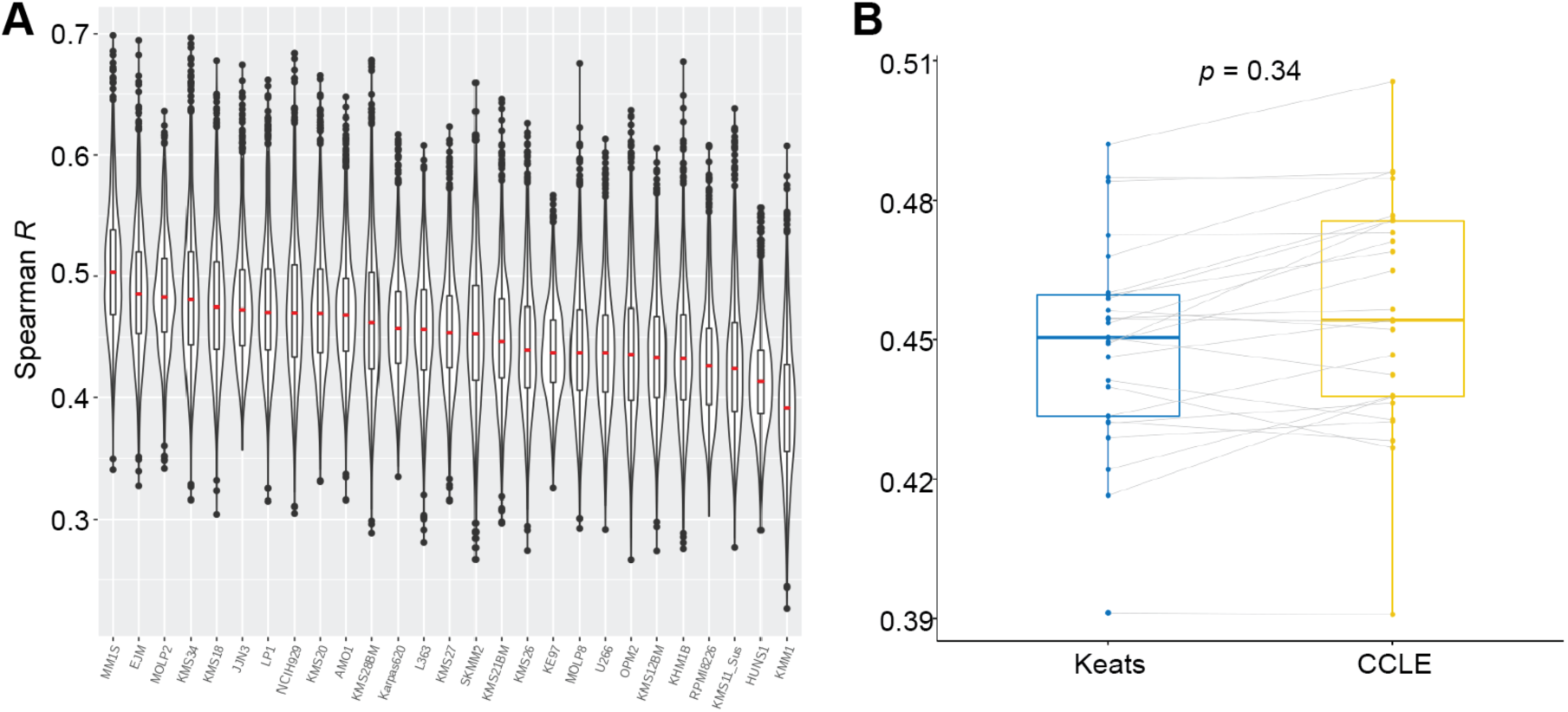
Ranking cell line similarity to patient tumor in CoMMpass using CCLE transcriptome data. **A.** Similar to Fig. 3A, correlation analysis of the CCLE and CoMMpass transcriptome data. Each sample in the violin plot corresponds to the Spearman correlation between one cell line and one primary tumor sample using the 5000 most variable genes. In the overlaid boxplot, the red center line depicts the median, the box limits depict the upper and lower quartiles, and the whiskers depict 1.5 times the interquartile range. **B.** Comparison of cell line ranking for the 25 lines included in both the CCLE and Keats datasets demonstrates no significant change in median correlation for each line as measured using each cell line dataset (*p*-value by Wilcoxon test).

**Figure S4.**
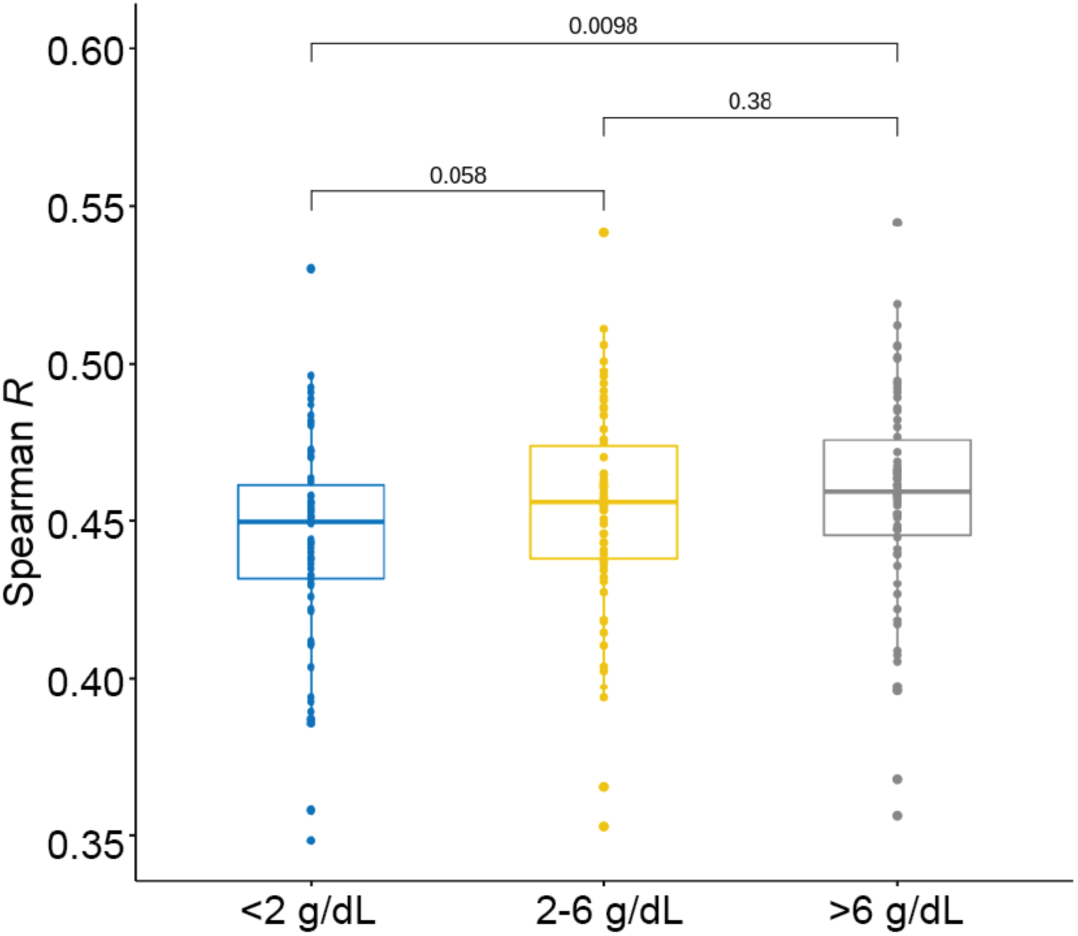
Patient M-spike at diagnosis correlates with increased similarity to cell lines. Box plots indicate median Spearman R for each cell line (dots) versus all patients as annotated in CoMMpass as having the indicated M-spike at diagnosis. Patient tumors associated with M-spike >6 g/dL have significantly greater similarity to cell lines than those with <2 g/dL. *p*-values by Wilcoxon test.

**Figure S5.**
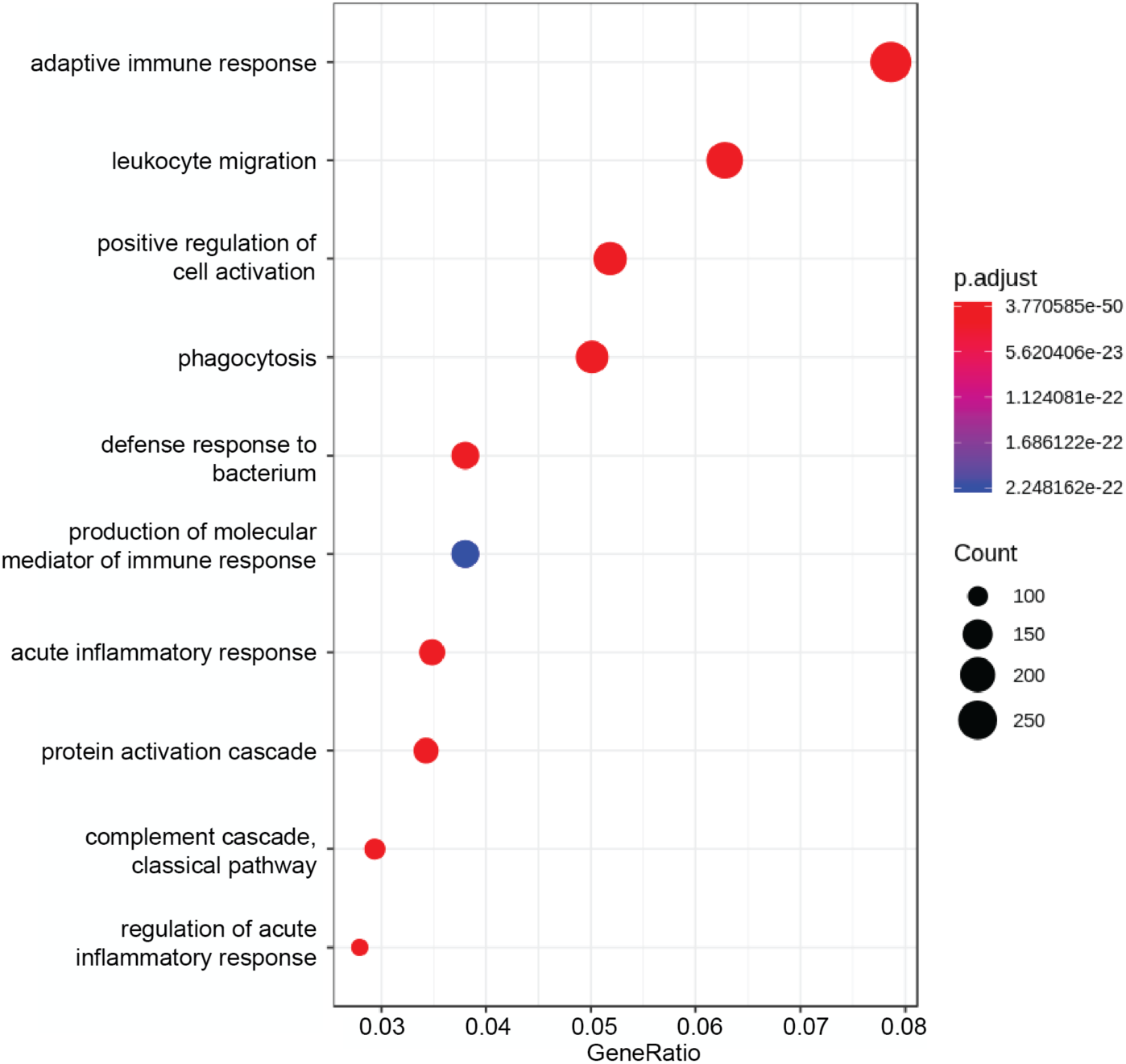
Gene Ontology Analysis recapitulates same upregulated biological functions in patient tumor vs. cell lines as found by GSEA (Fig. 5D).

**Figure S6.**
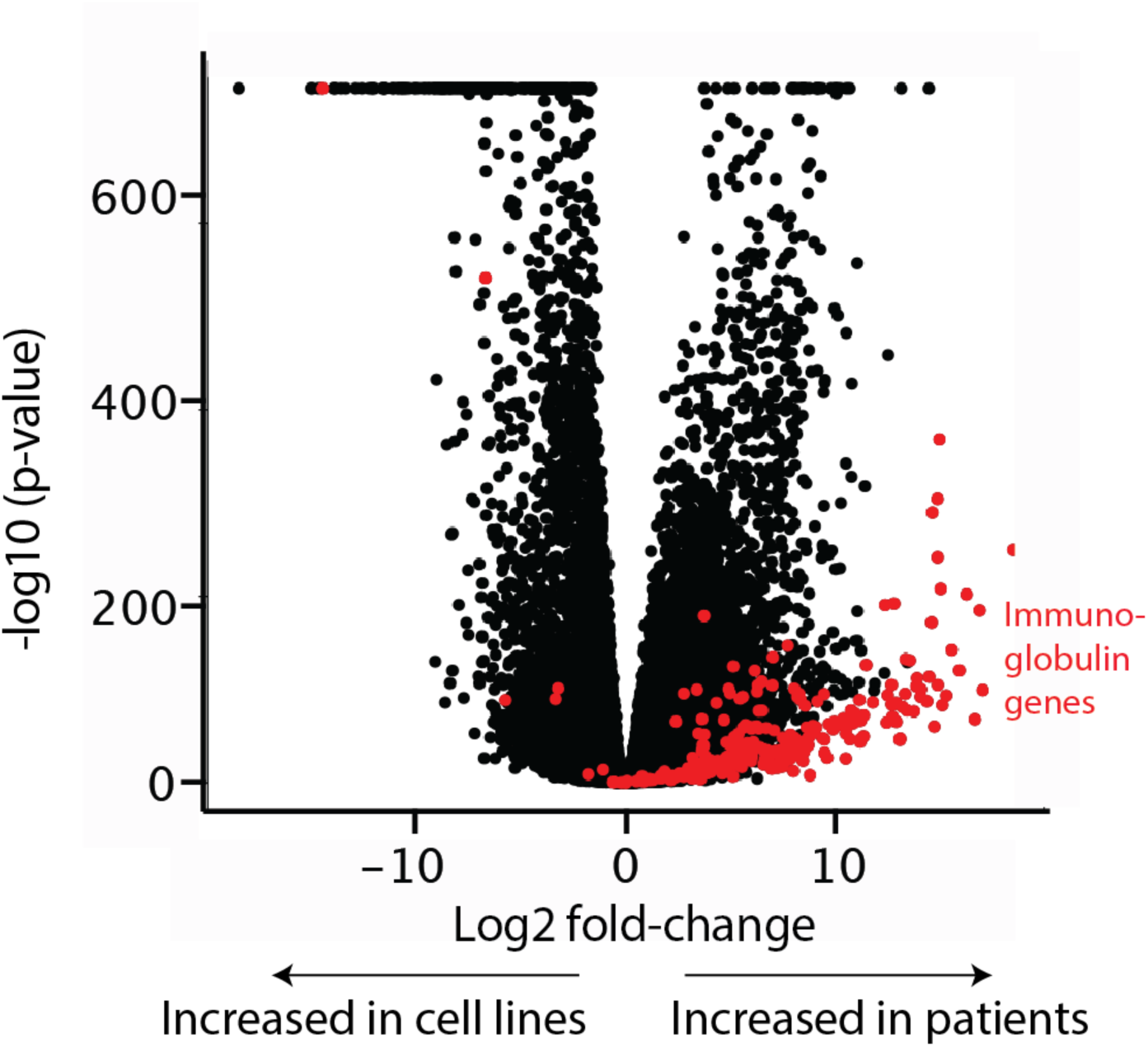
Overall aggregate comparison of gene-level expression between all patients and cell lines. Immunoglobulin genes, encoding different heavy and light chain isoforms (gene names beginning with “IGH”, “IGL”, and “IGK”), are highly represented among genes with increased expression in patients versus cell lines (also see Dataset S3). Comparison made using EdgeR tool on RNA-seq data.

**Figure S7.**
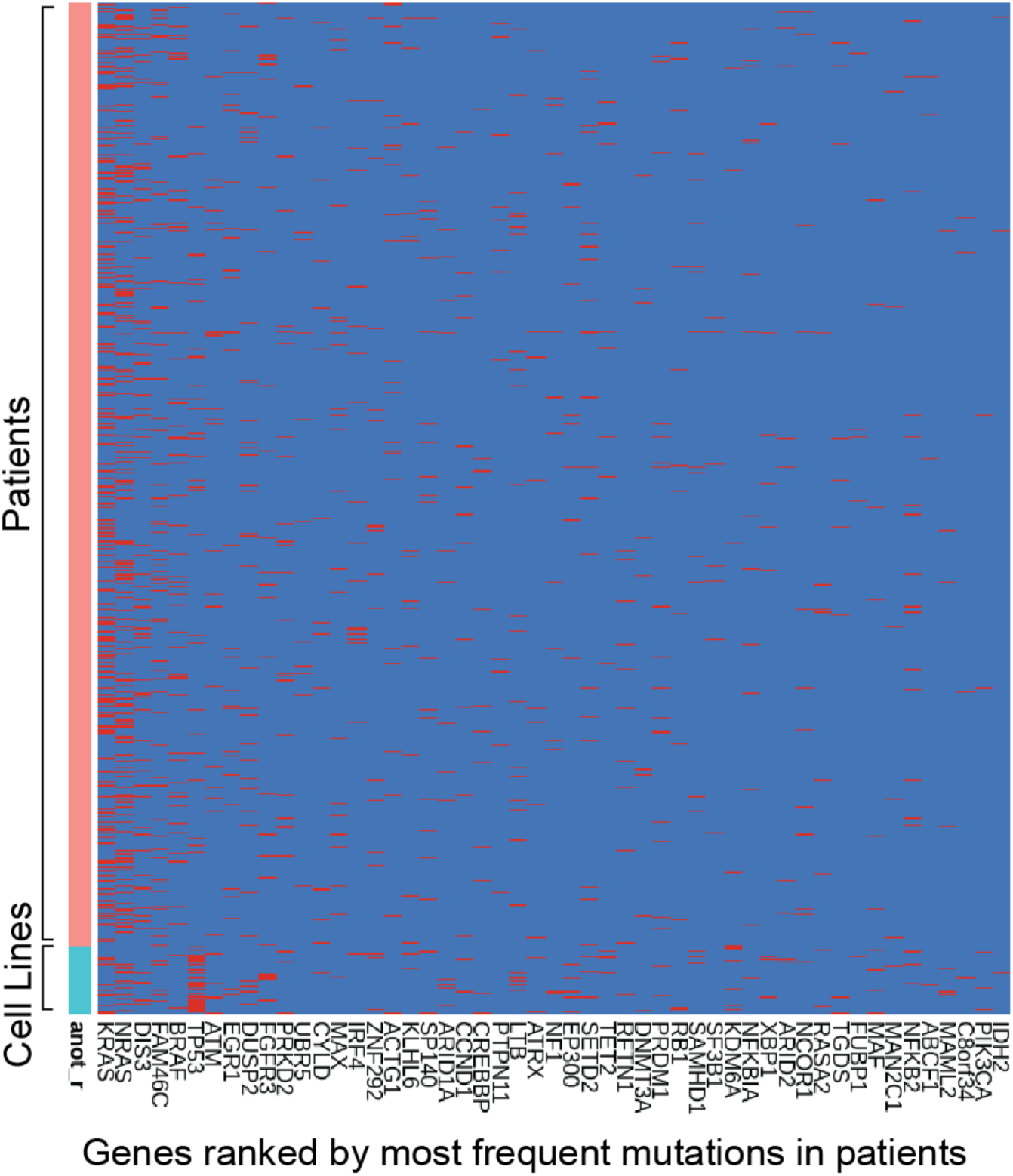
Mutation frequency in both cell lines and patients for most-common mutations detected in patient samples. Patient tumor mutation frequency as characterized by Walker *et al.* (*Blood*, 2018) and ordered by most to least frequent in patient samples. Red = mutation detected compared to hg19 reference genome. Blue = no mutation.

**Figure S8.**
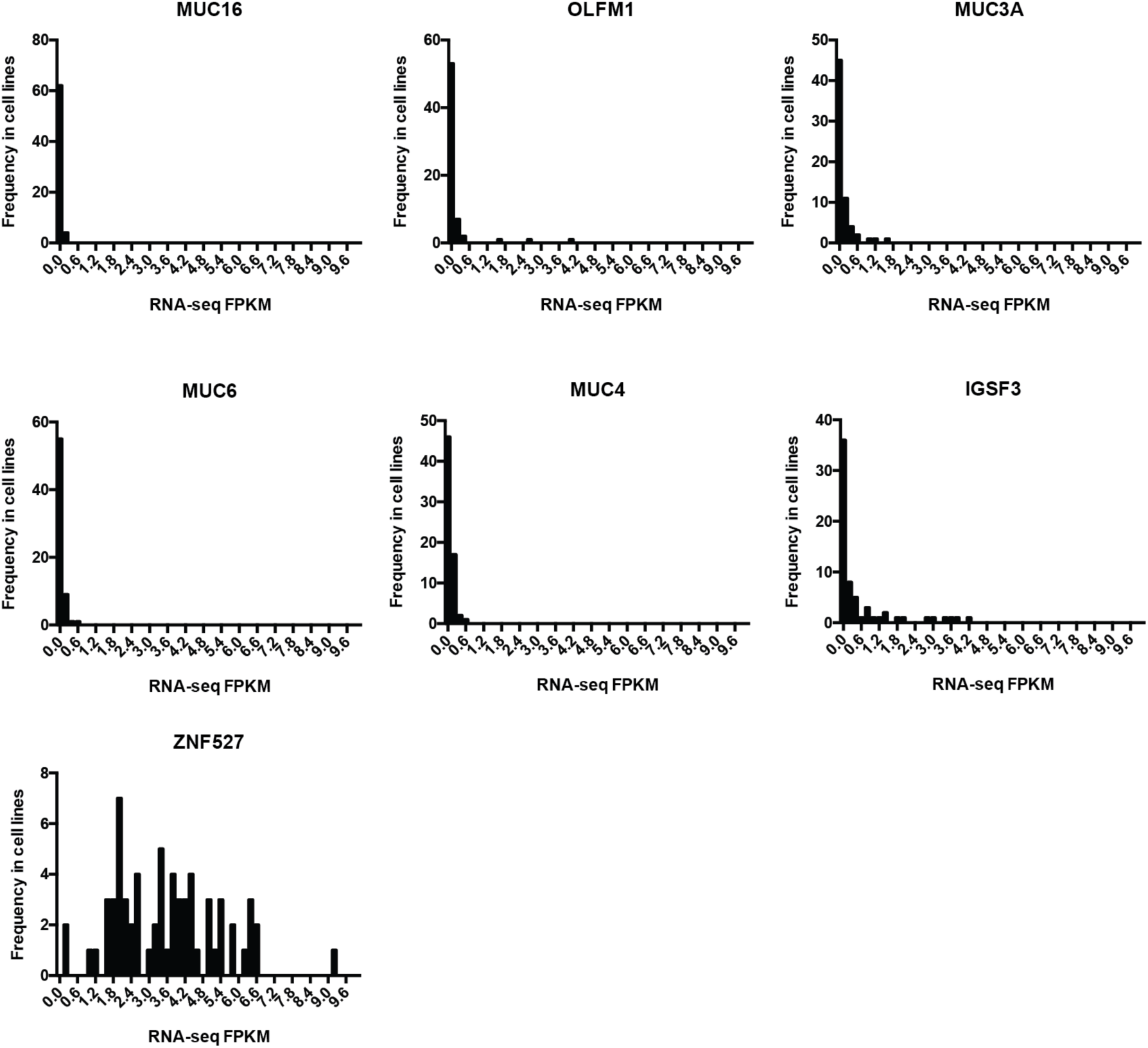
Expression levels of genes found to be frequently mutated in cell lines but not patients. Variant calls from processed exome sequencing data and processed transcript expression data, in FPKM, obtained from Keatslab.org.

**Figure S9.**
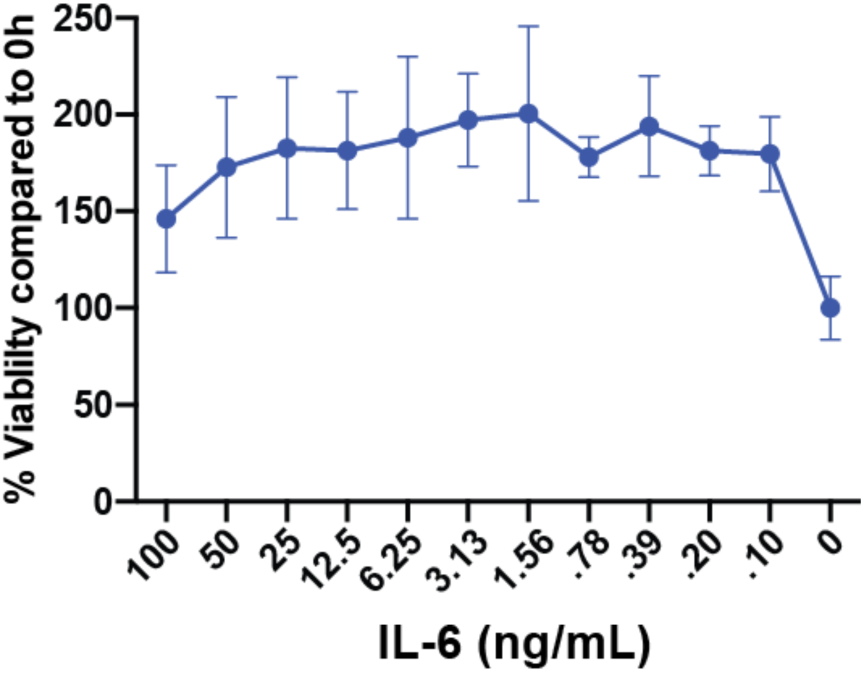
Proliferation of ANBL-6 across IL-6 concentrations. Our results confirm that luciferase-labeled ANBL-6 proliferates *in vitro* at IL-6 concentrations as low as 0.1 ng/mL (*n* = 4 per concentration; CellTiterGlo measured at 72h), as compared to baseline at 0h. No proliferation is observed without IL-6 (i.e. same number of cells present at 72h as at 0h).

## Supplementary Dataset Legends

**Supplementary Dataset 1 (.xlsx file):** *Sheet 1:* Data summary for patient-cell line transcriptional Spearman correlations including mean, median, and interquartile ranges. *Sheet 2:* Rankings of citations of each cell line per Google Scholar search. *Sheet 3:* Data summary for patient-cell line transcriptional Spearman correlations versus 25 overlapping myeloma cell lines in CCLE.

**Supplementary Dataset 2 (.xlsx file):** Cell line mutations found in exome sequencing data with variants called by Keats lab repository. Each sheet corresponds to each gene described in main text.

**Supplementary Dataset 3 (.xlsx file):** Differential expression data (log2-fold change and *p*-value) from EdgeR analysis for all genes as compared between cell lines and patient samples, and displayed in Fig. S6.

